# The RS Domain of Human CFIm68 Plays a Key Role in Selection Between Alternative Sites of Pre-mRNA Cleavage and Polyadenylation

**DOI:** 10.1101/177980

**Authors:** Jessica G. Hardy, Michael Tellier, Shona Murphy, Chris J. Norbury

## Abstract

Many eukaryotic protein-coding genes give rise to alternative mRNA isoforms with identical protein-coding capacities but which differ in the extents of their 3´ untranslated regions (3´UTRs), due to the usage of alternative sites of pre-mRNA cleavage and polyadenylation. By governing the presence of regulatory 3´UTR sequences, this type of alternative polyadenylation (APA) can significantly influence the stability, localisation and translation efficiency of mRNA. Though a variety of molecular mechanisms for APA have been proposed, previous studies have identified a pivotal role for the multi-subunit cleavage factor I (CFIm) in this process in mammals. Here we show that, in line with previous reports, depletion of the CFIm 68 kDa subunit (CFIm68) by CRISPR/Cas9-mediated gene disruption in HEK293 cells leads to a shift towards the use of promoter-proximal poly(A) sites. Using these cells as the basis for a complementation assay, we show that CFIm68 lacking its arginine/serine-rich (RS) domain retains the ability to form a nuclear complex with other CFIm subunits, but selectively lacks the capacity to restore polyadenylation at promoter-distal sites. In addition, nanoparticle-mediated analysis indicates that the RS domain is extensively phosphorylated in vivo. Overall, these results suggest that the CFIm68 RS domain makes a key regulatory contribution to APA.

## Introduction

Endonucleolytic cleavage of a eukaryotic pre-mRNA and subsequent polyadenylation defines the 3´ extent of the mature mRNA, and hence the extent to which its 3´ untranslated region (3´UTR) includes regulatory elements such as microRNA target sites and cis-acting sequences governing intracellular transport. Genome-wide transcript mapping studies suggest that more than half of all mammalian genes have multiple cleavage/polyadenylation sites (poly(A) sites) (1, 2), suggesting the potential for regulation of gene function at the level of alternative polyadenylation (APA) of the pre-mRNA. Indeed, for some genes subject to APA the intracellular location of protein synthesis has been shown to be determined by poly(A) site selection (3). For other genes, such as *CCND1*, which encodes the cancer-critical protein cyclin D1, use of a promoter-proximal, rather than promoter-distal, poly(A) site confers increased transcript stability and is associated with poor clinical outcomes (4-6). Indeed, transcriptome-wide shifts to favour the use of promoter-proximal poly(A) sites by hundreds of genes were found to accompany the onset of cell proliferation in T-lymphocytes and fibroblasts (7, 8), and to occur in a number of cancer cell lines compared with their normal counterparts (6).

The biochemistry of pre-mRNA cleavage and polyadenylation (CPA) is complex, with around 85 proteins having been identified in a purified 3’end processing complex in mammalian cells (9, 10). Among these, five core factors are necessary and sufficient to reconstitute efficient CPA in vitro (11). Four of these are multi-subunit complexes that together mediate the cleavage step - namely, cleavage and polyadenylation specificity factor (CPSF), cleavage stimulation factor (CstF), cleavage factor I (CFIm) and cleavage factor II (CFIIm). The fifth is the monomeric poly(A) polymerase (PAP) enzyme, which is recruited by interaction with CPSF and catalyses the polyadenylation reaction as well as contributing to the cleavage step. Productive CPA requires assembly of these core factors around the destined poly(A) site, which is partly mediated by recognition of cis-elements on the RNA. The CPSF complex recognises the conserved hexameric polyadenylation signal (PAS) of consensus sequence A(A/U)UAAA, which is located around 10-30 nucleotides (nt) upstream of the poly(A) site (12-15), while the CstF complex binds a more degenerate downstream GU-rich sequence (16-18). The roles of the CFIm and CFIIm complexes in the CPA reaction are less well defined, though CFIm is known to bind 40-50 nt upstream of poly(A) sites and may enhance CPSF recruitment (19). Assembly of the cleavage machinery stimulates enzymatic cleavage by the CPSF-73 subunit of CPSF (20) at a site between the PAS and the GU-rich sequence, following which PAP catalyses poly(A) tail addition.

While these details of the CPA reaction are relatively well established, it remains unclear how the choice between alternative poly(A) sites is regulated, and how such regulation changes under different physiological conditions. Many CPA factors and other RNA-associated factors have been implicated in poly(A) site selection, including CstF-64 (17), PABPN1 (21, 22), CPEB (23) and U1 snRNP (24). However, arguably the factor that has been most consistently and strikingly linked with APA is the cleavage factor I complex (CFIm) (reviewed in (25)).

CFIm is a metazoan-specific heterotetrameric complex composed of a homodimer of two 25 kDa subunits (CFIm25, also known as CPSF5, gene name *NUDT21*) and two larger subunits of either 59 kDa (CFIm59 or CPSF7) or 68 kDa (CFIm68 or CPSF6). It is thought a functional complex could contain one each of CFIm59 and CFIm68, or alternatively two CFIm59 or two CFIm68 subunits (26). Despite a lack of canonical RNA-binding motifs, CFIm25 is the major RNA-binding subunit of CFIm, forming sequence-specific interactions with UGUA sequences (27-30). This helps to position CFIm around 40-50 nt upstream of poly(A) sites, although it appears binding at this position can also occur in the absence of UGUA sequences (18). CFIm59 and CFIm68 are highly related in sequence and domain structure. Each has an RNA recognition motif (RRM), which rather than binding RNA forms an interaction surface with CFIm25, as well as a proline-rich region and a C-terminal RS domain, which in CFIm68 mediates interactions with splicing-related SR proteins (27).

Numerous studies have demonstrated that depletion of CFIm25 or CFIm68 from human cells leads to widespread 3´UTR shortening through increased relative usage of more proximal poly(A) sites (26, 31-33). The large number of affected transcripts and the overwhelming directionality of this effect suggests that CFIm may be a key regulator of APA, but a full understanding of its mechanism of action is lacking. CFIm may promote the use of distal poly(A) sites (18, 34), potentially through stabilising CPSF recruitment (19), but it has also been suggested that CFIm may alternatively or simultaneously repress the use of promoter-proximal poly(A) sites (35). Moreover, the broad physiological relevance of the CFIm knockdown phenotype is unclear. Aside from two examples of APA changes in neurological disease linked to altered CFIm expression (33, 36), there has been no demonstration that regulated changes in CFIm expression or activity orchestrate global changes in APA programmes.

Here, in order to investigate more fully the function of CFIm in APA in vivo, we used CRISPR/Cas9-based gene editing to generate a human cell line with substantial and permanent CFIm68 depletion, which demonstrated the expected 3´UTR shortening in representative transcripts. This cell line was used as the basis for a complementation assay to investigate the function of different CFIm68 mutant isoforms, which led to the discovery that CFIm68 variants lacking portions of the arginine/serine-rich (RS) domain show functional defects in APA regulation. In addition, western blot-based phosphorylation analysis identified multiple sites of serine phosphorylation within the RS domain. Our findings suggest that the RS domain makes a key contribution to the function of CFIm68 as a cleavage factor.

## Results

### Development of an APA complementation assay allowing analysis of CFIm68 function

In order to investigate the functions of specific domains and other sites of interest in CFIm68, we designed a complementation assay based on the well-defined 3´UTR shortening observable upon depletion of CFIm68 from human cells. Such an APA shift should be theoretically reversible by re-expression of wild type CFIm68 (complementation), but not by functionally impaired CFIm68 mutants. As a starting point, we attempted to completely abolish expression of CFIm68 in human embryonic kidney 293 (HEK293 Flp-In T-REx) cells using CRISPR/Cas9 genome editing (37), with a single guide RNA (sgRNA) targeted to the end of exon 3 of the *CPSF6* gene. Following puromycin enrichment and clonal isolation of cells transfected with the Cas9/sgRNA-expressing plasmid, anti-CFIm68 western blot screening identified several clones with decreased CFIm68 expression, confirming successful gene editing. Two cell lines were identified that showed CFIm68 expression at only ∼10% of the level seen in the parental cell line, and were named 68KD and 68KD-2 (Fig. 1A). From 43 CFIm68-targeted clones screened, none was identified with a total loss of CFIm68, and re-targeting of 68KD with a different sgRNA did not lead to generation of a complete knockout cell line (data not shown). This may suggest that CFIm68 is essential, although this is inconsistent with a previous report of CFIm68 knockout in similar cells (38).

**Figure 1.**
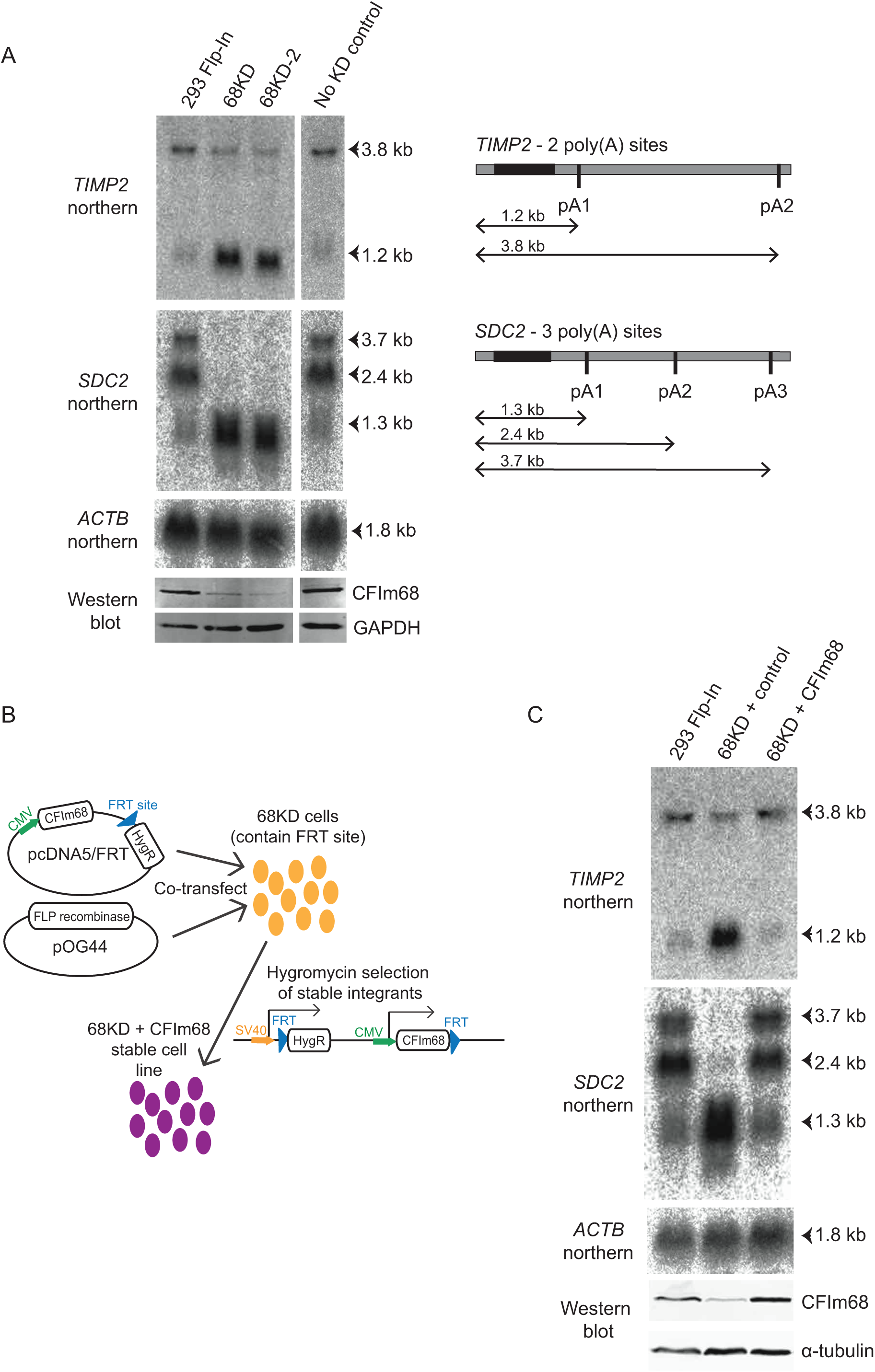
Establishment of a complementation assay based on APA regulation by CFIm68. A, Total RNA from CFIm68-depleted cell lines (68KD and 68KD-2) was analysed by northern blotting. Probes against the coding region of the TIMP2 and SDC2 genes allowed investigation of altered APA patterns by comparison with the parental 293 Flp-In cells, with positions of alternative poly(A) sites (pA) in these genes illustrated on the right. The ‘No KD’ control underwent CRISPR targeting but did not show CFIm68 depletion. ACTB acts as a loading control not subject to detectable APA. Western blotting (bottom) shows corresponding CFIm68 expression levels, with GAPDH as a loading control. B, An illustration of the Flp recombinase-mediated approach used to stably re-express CFIm68 in the 68KD cell line. C, Northern blotting/western blotting analysis of APA as described in A for stable cell lines with an empty plasmid (control) or CFIm68 plasmid integrated in the 68KD background.

Although complete knockout was not achieved, the 68KD and 68KD-2 cell lines with ∼90% depletion of CFIm68 were considered good candidates for further analysis. As a first step towards characterising these cells, the edited locus was amplified from purified genomic DNA and sequenced in order to identify the sequence changes leading to decreased protein expression. The results (Table 1) confirmed that both lines had at least one allele with only a small in-frame deletion, which would be expected to produce protein and probably therefore accounts for the faint CFIm68 band remaining on the western blot. All other alleles sequenced had newly-generated premature stop codons and would therefore be expected to undergo nonsense-mediated decay and not to produce functional protein. Only one such allele was sequenced for 68KD-2, whereas for 68KD, two premature stop alleles were identified. It is unclear whether we achieved full allele coverage even in 68KD, due to the karyotypic instability of HEK293 cells and resulting uncertainty surrounding *CPSF6* copy number. However, these results provide at least a partial view of the gene editing events underlying the observed CFIm68 protein depletion.

**Table 1:** *CPSF6* allele genotypes in CRISPR/Cas9-targeted CFIm68 knockdown cell lines.

| **Cell Line** | **CPSF6 allele 1** | **CPSF6 allele 2** | **CPSF6 allele 3** |
| --- | --- | --- | --- |
| 68KD | 3 bp deletion. Loss of K124. Presumably functional protein. | 18 bp deletion. Loss of S123, K124, G125 and splice donor site. Presumed premature stop codon inclusion through intron retention. | 8 bp deletion. Loss of K124, G125 and splice donor site. Presumed premature stop codon inclusion through intron retention. |
| 68KD-2 | 9 bp deletion. Loss of S123, K124, G125. Presumably functional protein. | 166 bp insertion. Generation of premature stop codon. | Not identified |

Based on previous reports (26, 32), we expected that the CFIm68 depletion in these cell lines would lead to general 3´UTR shortening compared to the parental cell line. To assess this, we performed northern blotting on isolated RNA from 68KD and 68KD-2 using DNA probes against the coding regions of two genes with well-defined APA sites – Tissue Inhibitor of Metalloproteinases 2 (*TIMP2*) and Syndecan 2 (*SDC2*) (31) (Fig. 1A, right panel). For both genes a clear change in cleavage site use was visible in both 68KD and 68KD-2 compared to the parental line, with a substantial increase in use of the most proximal poly(A) site and decreased use of more distal sites (Fig. 1A). This change was not observed in a control line, which had undergone the same CRISPR/Cas9 targeting procedure but had not been depleted of CFIm68, confirming that the APA shift was not a non-specific consequence of CRISPR/Cas9 targeting or cell cloning. As the 68KD and 68KD-2 lines showed the same phenotype, 68KD was considered representative and used for further experiments.

Phenotypic analysis of the 68KD cell line revealed a pronounced slow growth phenotype, with almost a three-fold increase in doubling time compared to its parental counterpart (Fig. 2A). This is consistent with a previous report (39), and suggests that CFIm68 depletion reduces cell fitness. In addition, a ChIP-sequencing analysis of global RNA polymerase II (pol II) distribution highlighted a clear global decrease in pol II occupancy at transcription start sites in 68KD cells by comparison with the parental line, suggesting that CFIm68 depletion impacts upon transcription initiation or elongation (Fig. 2B). Perhaps surprisingly, this ChIP-sequencing analysis did not reveal any obvious global alteration in pol II distribution around poly(A) sites (data not shown).

**Figure 2:**
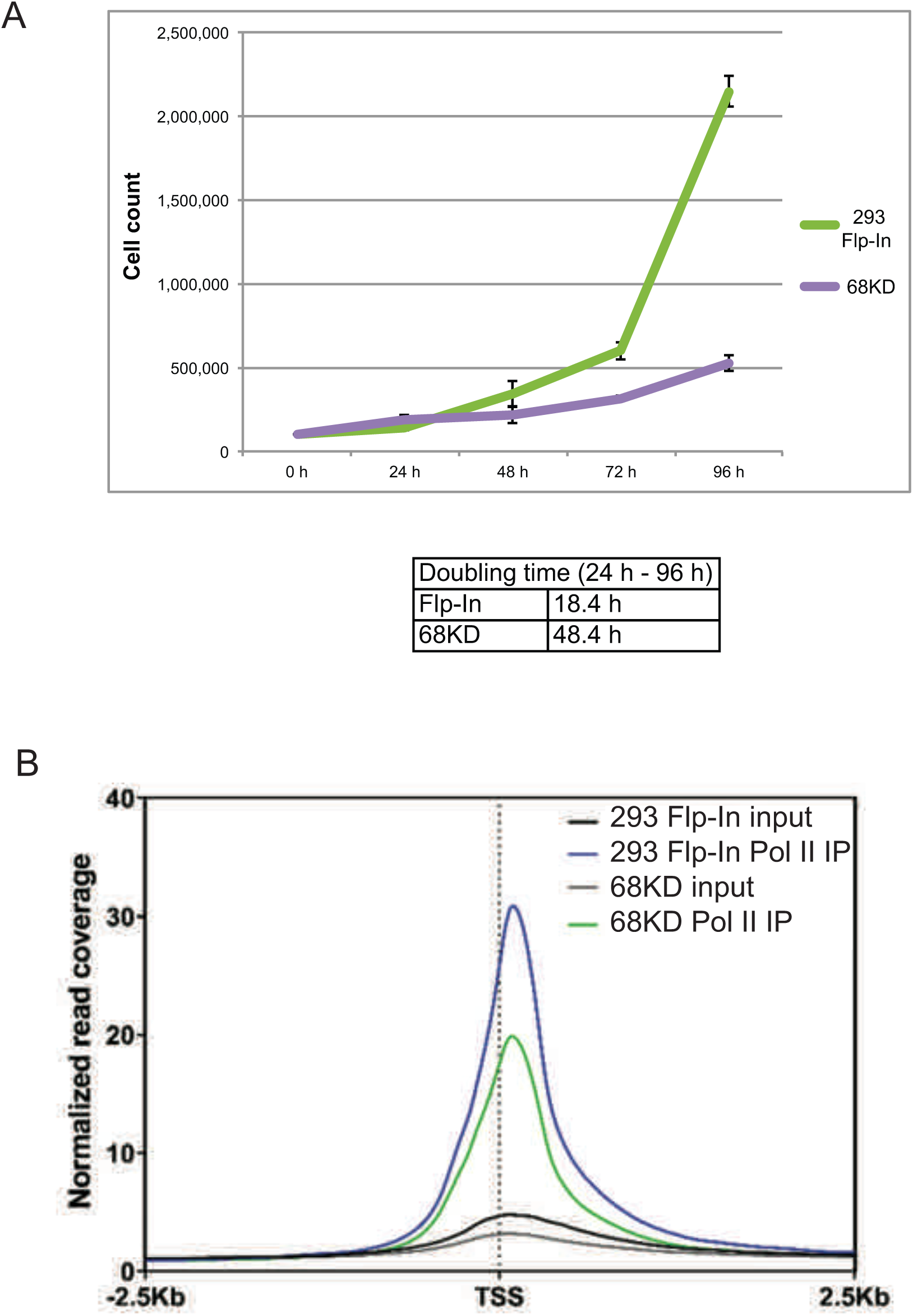
Reduced cell doubling rate and RNA polymerase II occupancy at the TSS in the 68KD cell line. A, Plots of mean cell counts across a 96 h timecourse in the 293 Flp-In and CFIm68-depleted 68KD cell line. Cells were seeded at 0 h in parallel wells of 100,000 cells each and a count was taken every 24 h. n=3, error bars represent ±S.E.M. B, Metaprofile of RNA polymerase II occupancy across 2200 protein-coding genes in the 293 Flp-In and 68KD cell lines, centred on the transcription start site (TSS), as measured in a ChIP-seq experiment.

To establish the complementation assay, we stably re-expressed CFIm68 in the 68KD cell line. This was achieved by exploiting the Flp recombinase target (FRT) site present in the parental cell line (HEK293 Flp-In T-REx), which facilitates genomic integration and stable expression of a gene of interest (Fig. 1B). Successful CFIm68 re-expression in the stable line (68KD + CFIm68) was confirmed by western blotting, which revealed a level of CFIm68 around 1.6-fold higher than in the parental cell line (Fig. 1C). *TIMP2*/*SDC2* northern blotting showed that this CFIm68 re-expression resulted in complete reversal of the 3´UTR shortening induced by CFIm68 knockdown (Fig. 1C), restoring the original APA profile observed in the parental cell line. The re-expression line was compared to a control line with an empty plasmid stably integrated (68KD + control), which as expected showed no restoration of CFIm68 expression and no APA complementation, confirming that this was a CFIm68-specific effect. Restoration of CFIm68 expression also reversed the growth defect observed in 68KD cells, confirming that the slow growth phenotype was a result of the CFIm68 depletion (data not shown).

While this report focuses on the CFIm68 complementation assay, it should be noted that we obtained similar results with a CFIm25-depleted line, which also showed a shift towards proximal poly(A) site use and provided the basis for a CFIm25 complementation assay (not shown).

### Interaction of CFIm68 with CFIm25 via its β2/β3 loop is required for normal APA regulation

Having established an effective APA complementation assay based on CFIm68 function, we investigated determinants of CFIm68 activity by comparing the complementation ability of mutant CFIm68 isoforms to that of wild type CFIm68. The first aspect of CFIm68 investigated was its interaction with CFIm25 within the CFIm complex, and whether this is required for regulation of poly(A) site selection by CFIm68. Residues 116-122 of the β2/β3 loop within the RRM of CFIm68 were first highlighted as likely mediators of interaction with CFIm25 based on crystal structural evidence (35), and have since been confirmed to be necessary for the CFIm25-CFIm68 interaction in human cells (39). We therefore generated a stable cell line to express CFIm68 Δ116-122 in the 68KD background (68KD + CFIm68 Δ116-122). Western blotting indicated that the CFIm68 Δ116-122 variant was expressed in this line at a level comparable to that of CFIm68 in the 68KD + CFIm68 cells (Fig. 3A, 3B).

**Figure 3.**
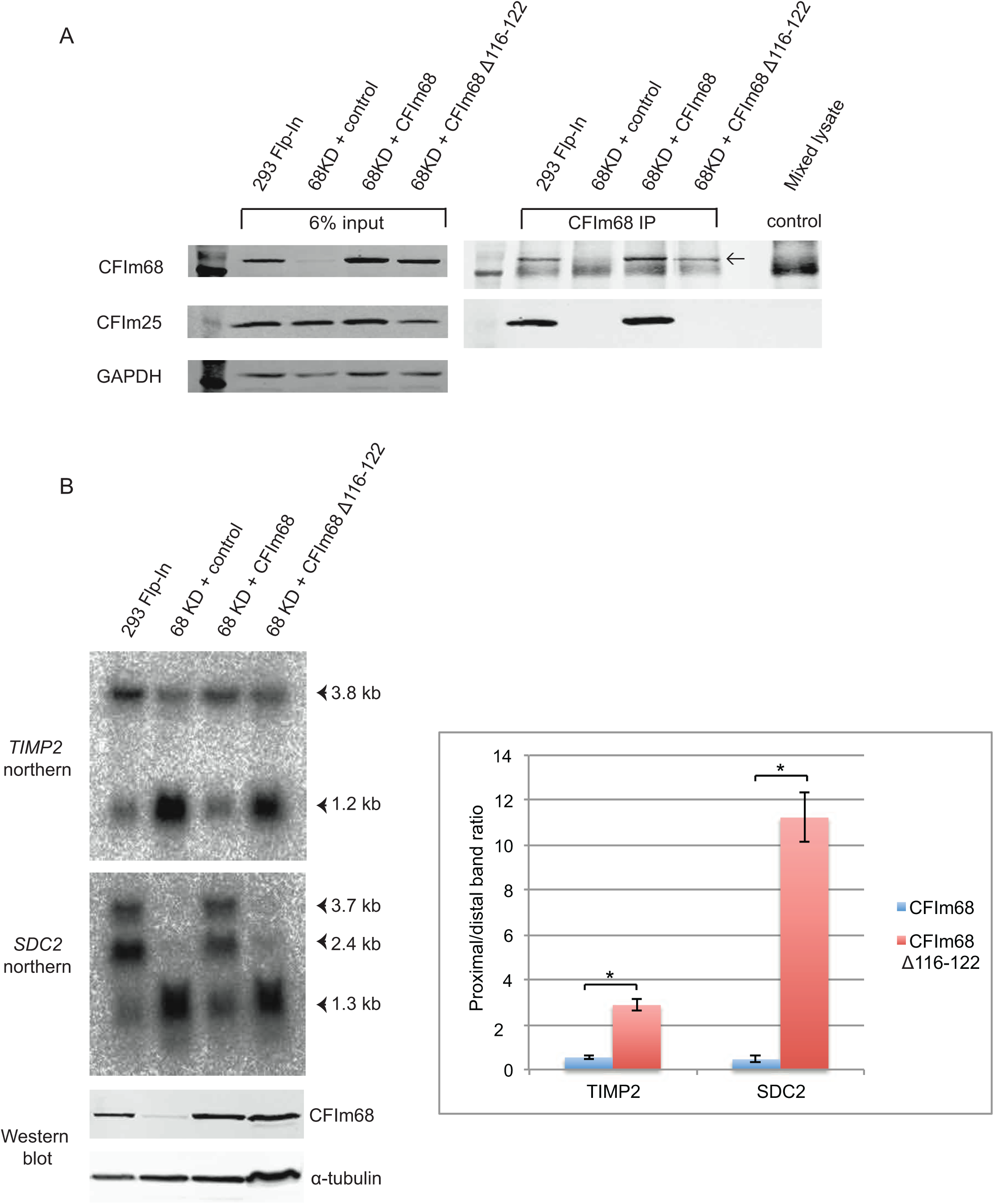
The CFIm68 Δ116-122 mutant is unable to interact with CFIm25 and cannot function in APA regulation. A, CFIm68 was immunoprecipitated from the given cell lines and western blotting was used to assess interaction with CFIm25. Whole cell lysate inputs are shown on the left with GAPDH as a loading control. IP samples are shown on the right, with the arrow highlighting the position of the CFIm68 band. The control IP was carried out on a mix of the 4 lysates using normal rabbit IgG. The result is representative of 2 biological repeats. B, Northern blotting was performed on total RNA from the given cell lines using probes against the TIMP2 and SDC2 genes as described in Figure 1, with corresponding CFIm68 western blots shown below (α-tubulin acts as a loading control). The graph on the right shows quantification of altered APA in the Δ116-122 mutant, expressed as proximal/distal ratios of band intensities measured by 2D densitometry (proximal = lower band, distal = sum of all other bands). n=2, error bars represent ± S.E.M. Statistical significance was calculated using a two-tailed student’s t-test.

Immunoprecipitation of CFIm68 Δ116-122 from this cell line confirmed that it was unable to interact detectably with CFIm25, whereas wild type CFIm68 from 68KD + CFIm68 cells immunoprecipitated CFIm25 efficiently (Fig. 3A). Having confirmed the lack of interaction, we used the complementation assay to investigate the functionality of the Δ116-122 mutant in poly(A) site selection. For both genes tested, there was a complete lack of complementation in the 68KD + CFIm68 Δ116-122 cell line, with the APA profile appearing similar to that of the 68KD + control line and showing a significant difference to that of the 68KD + CFIm68 line (Fig. 3B). This suggests that CFIm68 must interact with CFIm25 to regulate poly(A) site selection, and does not function in this process independently of the CFIm complex. The failure of the Δ116-122 mutant to complement APA despite its stable re-expression also validates the ability of the assay to detect functionally impaired CFIm68, demonstrating its suitability for use in investigating other determinants of CFIm68 activity.

### The RS domain of CFIm68 contributes to normal APA regulation

While the RRM of CFIm68 has a crucial role in APA regulation due to its involvement in the CFIm25-CFIm68 interaction, the contribution of the C-terminal arginine/serine-rich (RS) domain to APA is less clear. RS domains are found across the SR family of splicing regulators (40), and indeed the CFIm68 RS domain (Fig. 4A) has been shown to mediate interaction with other SR proteins such as 9G8 and SRp20 (27). A key role of this domain appears to be promoting nuclear import of CFIm68 via the transportin-3 (TNPO3) receptor (41), as well as promoting localisation to nuclear speckles (42). However, it has not been established whether the domain makes any contribution to the regulation of poly(A) site selection.

**Figure 4.**
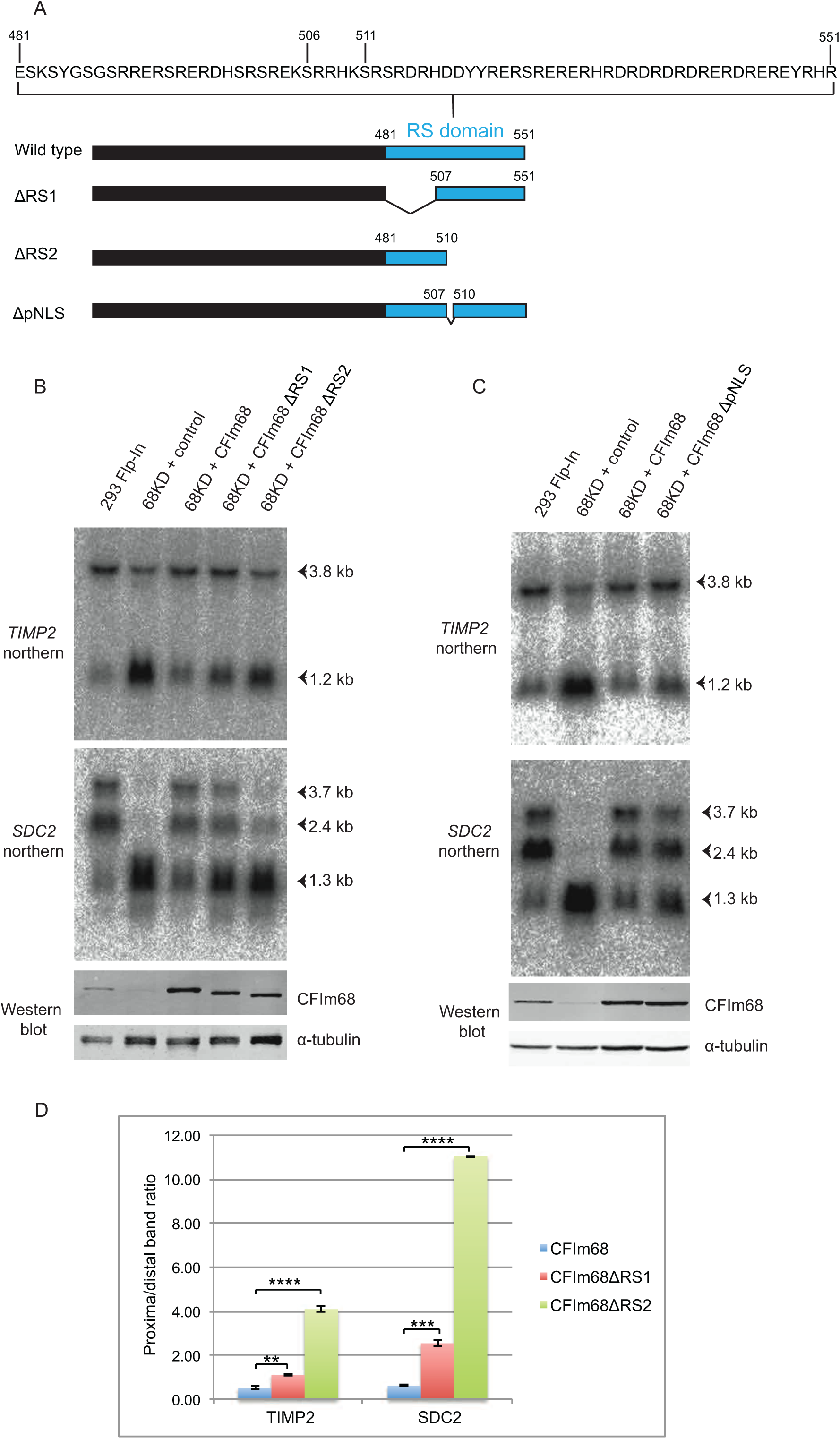
CFIm68 mutants lacking portions of the RS domain show impaired function in APA regulation. A, An illustration showing the position and sequence composition of the CFIm68 RS domain and the portions missing in the different RS deletion mutants. B/C, Northern blotting was performed on total RNA from the given cell lines using probes against the TIMP2 and SDC2 genes as described in Figure 1, with corresponding CFIm68 western blots shown below. Representative of 3 biological repeats (B) and 2 biological repeats (C). D, Quantification of altered APA in the ΔRS1 and ΔRS2 mutants, expressed as proximal/distal ratios of band intensities measured by 2D densitometry (proximal = lower band, distal = sum of all other bands). n=3, error bars represent ± S.E.M. Statistical significance was calculated using a two-tailed student’s t-test.

To investigate this, we generated stable cell lines expressing different CFIm68 RS domain deletion mutants. We aimed to maintain nuclear localisation of CFIm68, in order to probe for specific effects on poly(A) site selection rather than non-specific defects arising from mis-localisation. Two ‘half-domain’ deletion mutants (ΔRS1 and ΔRS2; Fig. 4A) were therefore generated, each removing a portion of the RS domain while leaving the remainder, including a putative nuclear localisation signal (NLS; residues 507-510, RRHK) intact. We also generated a line with deletion of this putative NLS alone (ΔpNLS; Fig. 4A). For unknown reasons, attempts to generate a line with the whole RS domain deleted were unsuccessful.

Despite robust expression of the ΔRS1 and ΔRS2 mutants, both showed impaired function in the complementation assay for both genes tested (Fig. 4B). This impairment was particularly marked for the ΔRS2 mutant, which showed an almost complete failure to restore distal APA to the *SDC2* gene. While the complementation defect for ΔRS1 was less pronounced, quantification of proximal/distal band ratios confirmed that both ΔRS1 and ΔRS2 showed significant differences to wild type CFIm68 for both genes (Fig. 4D). In contrast, the ΔpNLS mutant showed no obvious impairment in function (Fig. 4C). These results suggest that the RS domain, and particularly the region encompassing residues 511-551, makes a key contribution to the normal function of CFIm68 in poly(A) site selection.

The observation of this phenotype in the ΔRS cell lines raised the question of how loss of RS domain regions impairs CFIm68 activity. Although the available evidence indicates that the CFIm68 RRM domain is the key mediator of interaction with CFIm25 (27, 35), one possibility was that the ΔRS mutants are not incorporated into the CFIm complex effectively. However, immunoprecipitation of CFIm25 from the wild type and ΔRS cell lines confirmed that ΔRS1 and ΔRS2 were co-precipitated at levels comparable with those seen with wild type CFIm68 (Fig. 5A). This suggests that impaired CFIm complex formation is unlikely to explain the APA phenotype observed in the ΔRS mutant cell lines.

**Figure 5.**
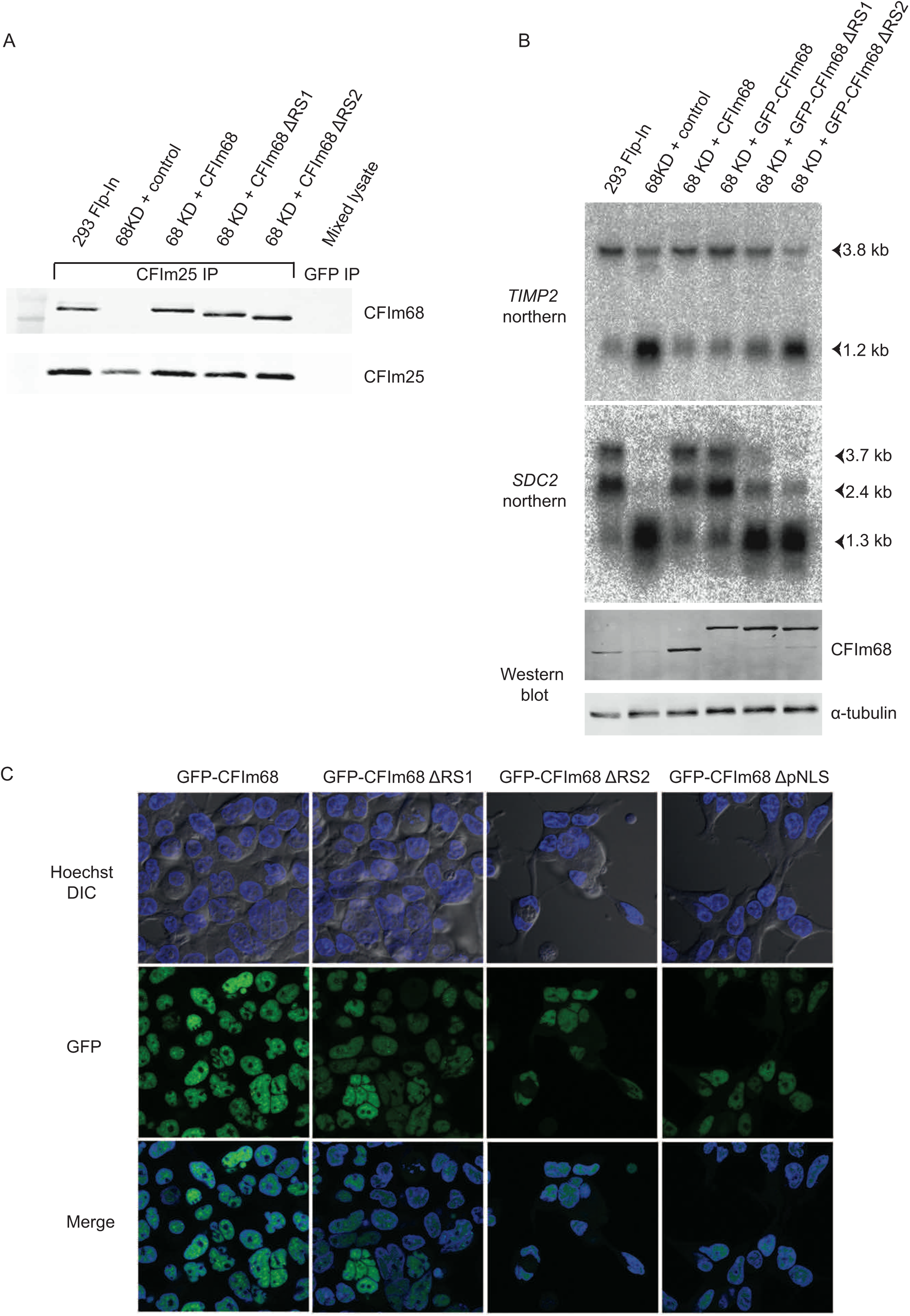
The CFIm68 ΔRS mutants are able to interact with CFIm25 and localise to the nucleus. A, CFIm25 was immunoprecipitated from the given cell lines and western blotting was used to assess interaction with CFIm68. The isotype control IP was carried out on a mix of the lysates using an anti-GFP antibody. B, Northern blotting was performed on total RNA from the given cell lines using probes against the TIMP2 and SDC2 genes as described in Figure 1, with corresponding CFIm68 western blots shown below. C, Live cell confocal microscopy was used to analyse the localisation of GFP-tagged CFIm68 isoforms in the given stable cell lines. Hoechst stain was added to the medium shortly before imaging to allow visualisation of the nucleus.

Another possible explanation for the APA phenotypes seen with the ΔRS mutants would be impaired nuclear import, as the RS domain has a reported role in this process (27, 41). To investigate this possibility, we generated cell lines expressing N-terminal GFP-tagged versions of the wild type and ΔRS CFIm68 isoforms. Analysis of APA in these cell lines using the complementation assay confirmed that GFP-CFIm68 could complement to the same extent as non-tagged CFIm68, suggesting that GFP tagging does not impair CFIm68 function (Fig. 5B). Consistently, GFP-ΔRS1 and GFP-ΔRS2 performed indistinguishably from their non-tagged counterparts (Fig. 5B; compare to Fig. 4B).

Live fluorescence imaging of these GFP-expressing cell lines was carried out, using a Hoechst counterstain to highlight nuclei. This revealed that wild type GFP-CFIm68 was clearly nuclear, as expected (Fig. 5C). Diffuse nuclear staining was observed, along with bright foci in many cells, consistent with previous reports of CFIm68 localisation to nuclear speckles and paraspeckles (42). GFP-ΔRS1 and GFP-ΔRS2 also showed clear nuclear localisation, albeit at a slightly lower intensity than wild type GFP-CFIm68 (Fig. 5C). This suggests that retaining either portion of the RS domain is sufficient to promote nuclear localisation, but it is not clear whether this requires the putative NLS, which is included in both mutants. Surprisingly, the GFP-ΔpNLS mutant retained nuclear localisation, suggesting that the RRHK sequence is dispensable for nuclear import (Fig. 5C). However, the intensity of signal was again lower than that seen with wild type CFIm68, and similar to that of the GFP-ΔRS1 and GFP-ΔRS2 mutants, suggesting that RRHK may function as a partial NLS in co-operation with the rest of the RS domain. Overall, these results suggest that the complementation defect observed with CFIm68 ΔRS mutants is not a simple consequence of failed nuclear localisation.

In summary, loss of portions of the CFIm68 RS domain, and in particular residues 511-551, leads to diminished CFIm68 function despite retention of nuclear localisation and CFIm complex formation. This suggests that the RS domain is required for the normal activity of CFIm68 in poly(A) site selection, and opens the door for further study into the contribution of the RS domain to CFIm68 function.

### Identification of serine phosphorylation sites in the CFIm68 RS domain

Having identified a key role for the CFIm68 RS domain in APA regulation, we were interested in the possibility that phosphorylation within this domain, or indeed elsewhere in the CFIm complex, may regulate CFIm activity to influence APA patterns. RS domains are known to be extensively serine phosphorylated in SR splicing factors, with phosphorylation/de-phosphorylation cycles playing a key role in regulating spliceosome assembly and splicing catalysis (43). Phosphorylation of the RS domain is not easily identifiable using standard tryptic peptide mass spectrometry, as the region has a high arginine density and is therefore digested into very short peptides by trypsin. Our attempts to overcome this limitation using an alternative protease, AspN, failed to give any coverage of the RS domain in mass spectrometry results, for unknown reasons.

An alternative methodology to probe CFIm phosphorylation was provided by the titanium-based nanoparticle reagent pIMAGO, which detects phosphorylation in a context-independent manner when applied to a western blot (44). We immunoprecipitated CFIm from HEK293 Flp-In cells using an anti-CFIm25 antibody and, following blotting, we used pIMAGO to detect phosphorylated proteins on the membrane. This approach yielded two clear bands on the blot, which corresponded exactly to the positions of CFIm68 and CFIm59 as identified by subsequent antibody detection (Fig. 6A). Furthermore, the upper band was absent from immunoprecipitates of extracts from the 68KD + control line and was restored in the 68KD + CFIm68 line, confirming that this band represented phosphorylated CFIm68 (Fig. 6A). The loss of the lower band in the 68KD + CFIm68 line corresponds with the concomitant reduction in CFIm59 levels (a result of CFIm68 overexpression), supporting the conclusion that this band represents phosphorylated CFIm59. Use of pIMAGO therefore showed that CFIm59 and CFIm68, but not CFIm25, are detectably phosphorylated in HEK 293 Flp-In cells.

**Figure 6.**
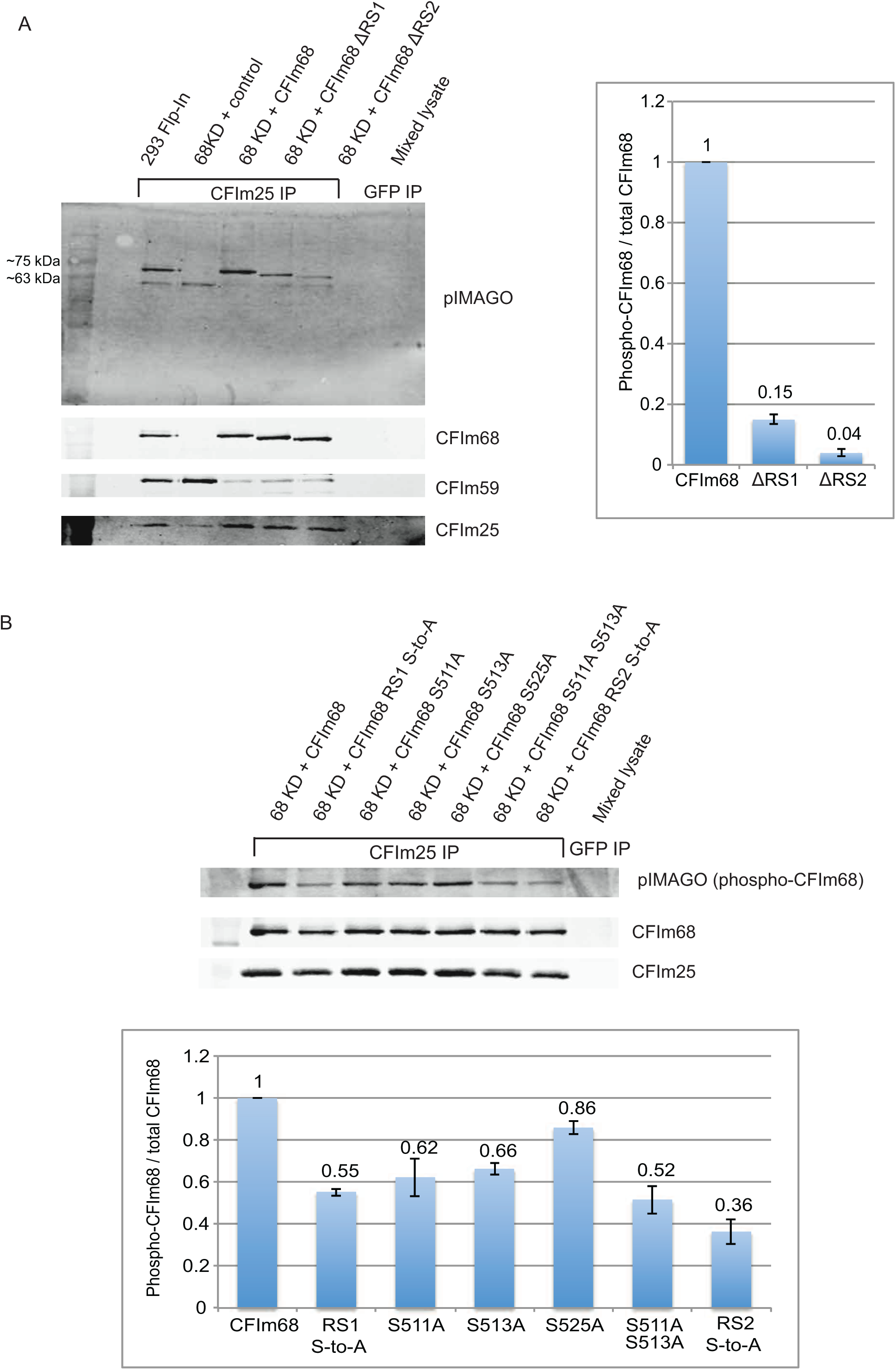
The CFIm68 RS domain is extensively serine-phosphorylated. CFIm25 was immunoprecipitated from the given cell lines and samples were analysed by western blotting. Anti-GFP IP was used as a negative isotype control. The nanoparticle pIMAGO reagent was used to detect phosphorylated proteins, followed by standard antibody-mediated detection of CFIm subunits. Fluorescent band intensities for phosphorylated and total CFIm68 were quantified and results were expressed as a ratio of phospho-CFIm68/total CFIm68, standardised to a value of 1 in the 68KD + CFIm68 cell line. n=3, error bars represent ± S.E.M. A, The effect of RS1 or RS2 region deletion on total phosphorylation level. B, The effect of serine-to-alanine mutagenesis within the RS1 and RS2 regions on total phosphorylation level.

When the same approach was applied to the ΔRS1 and ΔRS2 lines, there was a substantial decrease in the CFIm68 pIMAGO signal for both cell lines compared to the wild type, despite equivalent levels of total CFIm68 expression (Fig. 6A). Quantification of the pIMAGO signal normalized to total CFIm68 indicated an 85% decrease in phosphorylation for ΔRS1 and a 96% decrease for ΔRS2. This suggests that the majority of CFIm68 phosphorylation is found within the RS domain.

To determine which RS domain residues are phosphorylation targets, we performed site-directed mutagenesis. Initially, phospho-ablating serine-to-alanine (S-to-A) and tyrosine-to-phenylalanine (Y-to-F) mutations were generated at the six potential phosphorylation sites in the RS2 region, which appeared to be more heavily phosphorylated than the RS1 region. We generated stable cell lines expressing the phosphosite mutants and analysed these using the pIMAGO approach (Fig. 6B). S511A or S513A mutation led to a clear decrease in phosphorylation signal (38% and 34% decrease respectively), with a smaller decrease (14%) seen for S525A. The triple S511A S513A S525A (RS2 S-to-A) mutant showed a 64% loss of phosphorylation, suggesting that phosphorylation of these three residues together accounts for a substantial proportion of overall CFIm68 phosphorylation. The three Y-to-F mutations had no effect on phosphorylation (data not shown). Interestingly, the loss of phosphorylation seen for the RS2 S-to-A mutant (Fig. 6B) did not fully recapitulate the more substantial loss observed in the ΔRS2 line (Fig. 6A).

To investigate phosphorylation in the RS1 domain, we generated a mutant line with simultaneous S-to-A conversion of all eight serines in this region (RS1 S-to-A). pIMAGO analysis of this stable cell line revealed a 45% loss in phosphorylation signal, confirming that one or more of these serine residues is indeed phosphorylated (Fig. 6B). Mutation of the single tyrosine (Y485F) did not have any effect on phosphorylation (data not shown). Again, the drop in signal for RS1 S-to-A (Fig. 6B) did not recapitulate the loss seen on deletion of the RS1 region (ΔRS1; Fig. 6A), Interestingly, repeated attempts to generate a cell line expressing a full RS S-to-A mutant encompassing both the RS1 and RS2 regions were unsuccessful, suggesting that this mutant non-phosphorylatable isoform may have a lethal dominant negative effect.

Having identified extensive phosphorylation within the CFIm68 RS domain, we wanted to investigate the potential functional importance of these sites. We analysed RNA from various stable cell lines with the S-to-A mutations described above or with phospho-mimetic serine-to-aspartate (S-to-D) mutations in the northern blot complementation assay (Fig. 7). None of the single or combined RS2 domain mutants, whether S-to-A or S-to-D, showed any obviously altered complementation activity, suggesting that phosphorylation of S511/S513/S525 does not play a direct role in APA regulation. In addition, the RS1 S-to-A mutant not only complemented fully, but led to a small relative increase in use of distal poly(A) sites for both genes tested. For the mutants studied, however, there were no substantial impairments in complementation ability comparable to those seen in the ΔRS1 and ΔRS2 lines (Fig. 4B). Overall, these results suggest that serine phosphorylation in the RS domain of CFIm68 does not play a major role in regulating alternative polyadenylation.

**Figure 7.**
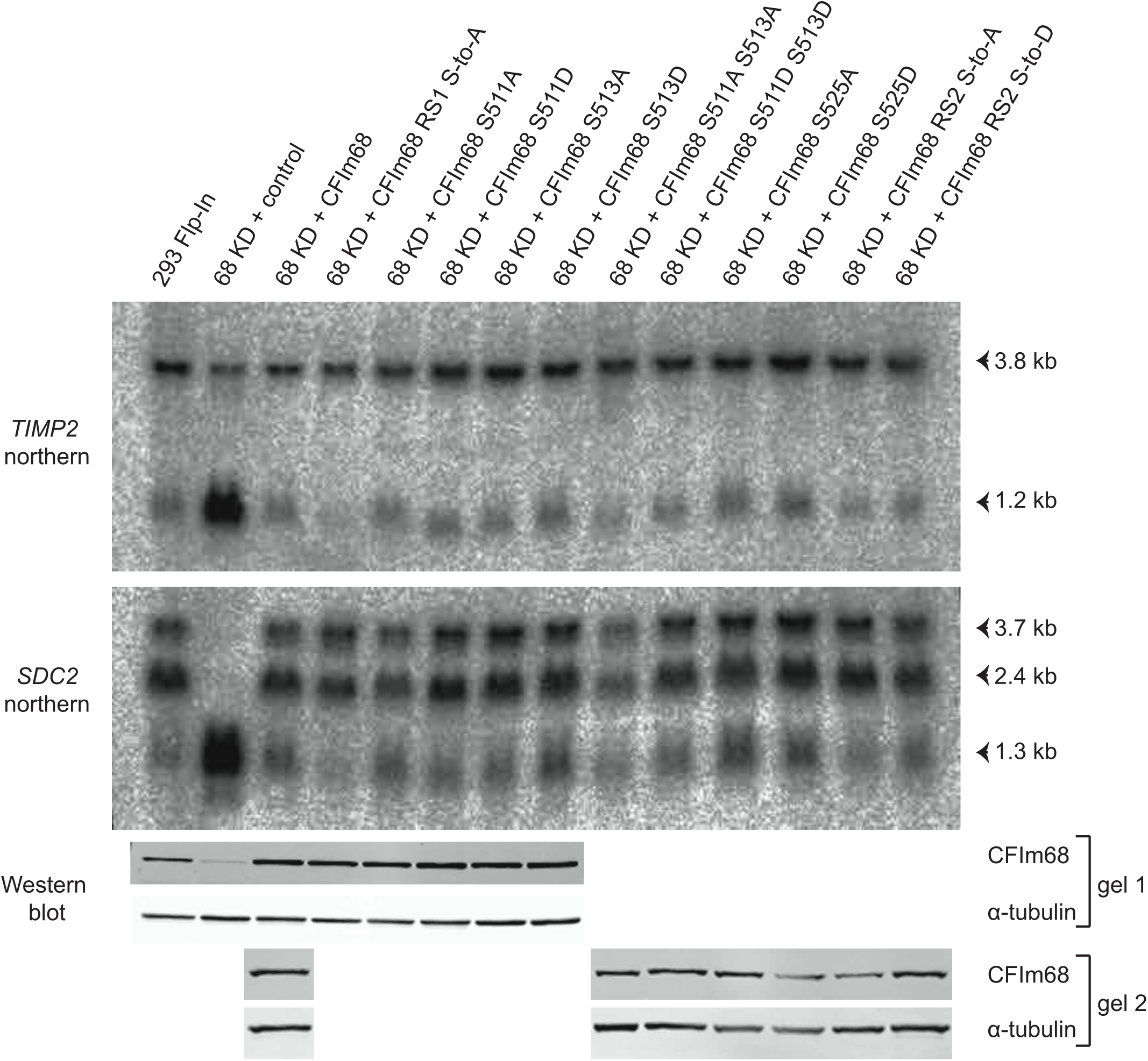
Phospho-ablating and phospho-mimetic serine mutagenesis in the CFIm68 RS domain does not alter APA regulation. Northern blotting was performed on total RNA from the given stable cell lines using probes against TIMP2 and SDC2 as described in Figure 1, with corresponding western blots shown below. Samples were run across two western blots as indicated, and wild type (68KD + CFIm68) sample was included on both gels to allow direct comparison with all mutants.

## Discussion

In this work, we established a complementation assay based on the activity of CFIm68 as an APA regulator and used it to probe various features of CFIm68 function. A clear and permanent shift towards increased proximal poly(A) site usage was achieved through CRISPR/Cas9-mediated CFIm68 depletion, and this was completely reversible by re-expression of CFIm68. By using the Flp-In system to provide stable and permanent re-expression of different CFIm68 isoforms from a common genomic locus, robust comparisons could be made between the complementation ability of these isoforms. This versatile assay could be exploited to examine other aspects of CFIm68 structure/function in addition to the domains and post-translational modification sites investigated here.

The finding that the CFIm68 RS domain makes a key contribution to APA regulation raises several questions, and we anticipate that further work will focus on the specific mechanistic roles of this domain. It was perhaps surprising that the GFP-CFIm68 ΔRS2 mutant showed predominantly nuclear localisation, given previous observations that a similar mutant with only additional deletion of the pNLS (RRHK, residues 507-510) showed a diffuse cytoplasmic/nuclear distribution (27). Our microscopy results suggest that this pNLS acts at most as a weak NLS, but it is certainly possible that its additional loss, when combined with loss of the RS2 region, leads to a substantial reduction in nuclear import. Ideally, the contribution of different regions of the RS domain to nuclear import would be more fully dissected through construction of a wider range of mutants, including the full ΔRS (Δ481-551) mutant, which we were unable to study here.

Further insight may also be gained from defining the interaction interface of the CFIm68 RS domain with the nuclear import factor transportin 3 (Tnpo3), which has been strongly implicated in CFIm68 nuclear import in the context of HIV infection (45). Mutational studies have identified the arginine-rich helix of Tnpo3 HEAT repeat 15 as important for the Tnpo3-CFIm68 interaction (41), but the CFIm68 residues involved have yet to be fully determined. Interaction of the same Tnpo3 arginine-rich helix with the SR protein ASF/SF2 absolutely requires serine phosphorylation within the RS domain, but this does not appear to be the case for CFIm68 (41, 46). It may be that the numerous phospho-mimetic residues within RD/RE dipeptides of the CFIm68 RS domain negate the serine phosphorylation requirement for nuclear import.

While the GFP-ΔRS1 and GFP-ΔRS2 mutants were clearly predominantly nuclear, the GFP intensity was slightly lower than that seen with wild type CFIm68. It could therefore be postulated that the lower level of ΔRS mutants reaching the nucleus by comparison with the wild type accounts for the impaired complementation phenotype. However, this seems unlikely, as despite the apparent similarity in GFP intensity between the ΔRS mutants, including ΔpNLS, ΔRS2 was substantially more functionally impaired than the others. Moreover, the fact that CFIm68 is overexpressed in these Flp-In cell lines would suggest that even with decreased nuclear import efficiency there may be a similar level of CFIm68 reaching the nucleus here as in parental Flp-In cells.

One more likely possibility is that altered subnuclear localisation of ΔRS mutants impairs their function. Previous reports have shown that the speckled distribution of CFIm68, which we also observed (Fig. 5), results from a combination of RRM-mediated localisation to paraspeckles and RS domain-mediated targeting to the periphery of nuclear speckles (42). Nuclear speckles, also defined as interchromatin granule clusters (IGCs), are subnuclear structures enriched in a range of RNA-associated factors, and in particular pre-mRNA splicing factors (reviewed in (47)). They are not thought to be sites of active transcription, but rather may act as storage or assembly compartments for RNA processing factors, which may be poised for recruitment to transcription sites. This raises the possibility that decreased accumulation of CFIm68 in nuclear speckles upon RS domain loss may decrease the efficiency with which it is recruited to transcribing genes, potentially contributing to the observed switch towards proximal poly(A) site use. Alternatively, RS domain-mediated interactions with other proteins within nuclear speckles, rather than the speckle localisation itself, may be key in regulating poly(A) site choice.

More generally, it may be that the contribution of the RS domain to promoting selection of distal poly(A) sites stems from interactions with specific proteins around these sites. There is substantial evidence that interactions between splicing and cleavage/poly(A) factors are crucial in definition of the 3´ terminal exon (48-50), and it is conceivable that such interactions between CFIm68 and other SR proteins may promote selection of distal poly(A) sites or inhibit selection of proximal poly(A) sites. RS domain-mediated interactions of CFIm68 with the SR proteins 9G8, TRA2β and SRp20 have already been identified (27), while the homologous CFIm59 forms an interaction with U2AF65 that stimulates 3´ end processing (51). The difference in interacting partners between the CFIm68 and CFIm59 SR domains is notable, and may partially account for their different functions – specifically, that CFIm59 does not play any obvious part in poly(A) site selection (26, 32).

Another possibility is that the CFIm68 RS domain contributes to poly(A) site selection through RNA binding. While CFIm25 is the major RNA-binding subunit of CFIm, the CFIm68 RS domain has measurable affinity for RNA in vitro (27). However, this may well be a result of non-specific ionic interaction, and it is unclear whether this represents an important role of the RS domain in vivo.

The finding that the RS domain is extensively serine phosphorylated is in line with the known phosphorylation of similar domains in other SR family proteins (reviewed in (43)). Site-directed mutagenesis coupled with pIMAGO western blot analysis allowed more detailed elucidation of the distribution of phosphorylation within the domain, revealing that both the RS1 and RS2 regions are phosphorylated and highlighting S511 and S513 as particularly highly-phosphorylated residues. An interesting observation was that full S-to-A mutagenesis within either RS1 or RS2 did not fully recapitulate the greater loss of phosphorylation observed upon deletion of the domains (ΔRS1/ΔRS2). This suggests that the RS1 and RS2 regions also have an indirect effect on overall CFIm68 phosphorylation in addition to being targets themselves. For example, the conformation and accessibility of the RS1 domain may be altered in the absence of RS2, perhaps impairing kinase recruitment and thus further decreasing overall phosphorylation levels. A fuller understanding of the phosphorylation dynamics and stoichiometry within this domain will likely require full characterisation of the upstream signalling pathways and kinases responsible. The SRPK and CLK family kinases represent strong candidates, as they have a known role in phosphorylating other SR family proteins with similar target motifs (reviewed in (43)). Indeed, the ability of SRPK1 to phosphorylate a heterologously expressed CFIm68 RS domain has been demonstrated in vitro (41).

The role of RS domain phosphorylation in regulating CFIm68 activity remains to be fully elucidated, and it was perhaps surprising that most of the phospho-ablating or phospho-mimetic mutants studied showed no obvious alteration in APA regulation. It is possible that there is some degree of redundancy in phosphorylation of this region, such that loss of some sites is compensated by retention of others. Interestingly, as for the full ΔRS mutant, attempts to generate a cell line expressing the full RS S-to-A mutant encompassing all serines in the RS1 and RS2 domains were unsuccessful, suggesting that complete loss of this phosphorylation may be selectively disadvantageous. In addition, the observation of relatively enhanced distal poly(A) site use in the RS1 S-to-A mutant warrants further study. It would be of considerable interest to analyse the phospho-mimetic RS1 S-to-D equivalent, to investigate the intriguing possibility that phosphorylation in this RS1 region may act to promote increased relative proximal poly(A) site selection by CFIm68.

If SR domain phosphorylation should ultimately prove dispensable for normal APA regulation, the question would remain of what role phosphorylation plays in CFIm68 biology. It is important to note the limitations of the complementation assay used here, which only looked at two model transcripts. Extension of APA analysis to the transcriptome-wide level may be necessary to reveal subtler, transcript-specific effects of SR domain phosphorylation. The potential ability of CFIm to regulate poly(A) site selection at a subset of transcripts in response to physiological stimuli remains an intriguing possibility, deserving of further study.

## Experimental Procedures

### Cell culture

HEK293 Flp-In T-REx cells (Invitrogen, catalogue no. R78007) were cultured in DMEM (Sigma) supplemented with 10% FBS (Gibco), 100 units/ml penicillin + 100 μg/ml streptomycin (Gibco), 15 μg/ml blasticidin (InvivoGen) and 100 μg/ml zeocin (InvivoGen).

Flp-In-derived stable cell lines were cultured in DMEM supplemented as described above, with 150 μg/ml hygromycin B (InvivoGen) replacing 100 μg/ml zeocin.

For cell growth curve measurements, cells were harvested from parallel wells by trypsinisation at 24 h intervals and counted using Glasstic slides with grid (KOVA) after diluting 5-fold in 0.4% trypan blue solution (Sigma).

### CRISPR/Cas9 gene editing

For Cas9 targeting of the *CPSF6* gene, forward and reverse oligonucleotides encompassing the sgRNA target sequence CGGGCAAATGGCCAGTCAAAGGG were annealed and inserted into the pSpCas9(BB)-2A-Puro (PX459) plasmid using *Bbs*I digest and ligation as described (37).

HEK293 Flp-In T-REx cells were seeded in a 35 mm well and transfected after 24 h with 2 μg of the resulting plasmid. Transfections were performed using the JetPrime reagent and protocol (Polyplus-transfection) with 2 μl JetPrime per μg DNA. After 24 h, 2.5 μg/ml puromycin was added to the medium, and after a further 24 h, cells were passaged into a 10 cm dish with 2.5 μg/ml puromycin.

72 h after transfection, cells were diluted to isolate single cells and obtain clonal populations, either by plating at low density in 10 cm dishes, or by serial dilution in 96-well plates. One to two weeks later, single colonies were identified, dissociated using TrypLE (Thermo Fisher) and expanded for analysis.

### Sequencing of the edited CPSF6 locus

Genomic DNA (gDNA) was purified from 68KD cells using the Wizard SV Genomic DNA Purification System (Promega). The sequence around the Cas9-targeted site in the *CPSF6* gene was amplified by PCR with Simpler Red Taq polymerase (ThermoScientific) from 100 ng gDNA using the ‘CFIm68 target site’ F/R oligonucleotides. Total or gel-purified PCR product was ligated into the pCR2.1 plasmid using a TA cloning kit (Invitrogen) and Sanger sequencing of individual inserts was performed using the M13 R oligonucleotide.

### Generation of stable Flp-In cell lines

For generation of the CFIm68 re-expression plasmid, the *CPSF6* coding sequence was originally amplified from a human fibroblast cDNA library (kindly provided by Hiroto Okayama) using *Not*I-CFIm68 F and CFIm68-*Xho*I R oligonucleotides. *Kpn*I/*Xho*I digest was ultimately used to excise the *CPSF6*-containing fragment from an intermediate plasmid and ligate it into pcDNA5/FRT (Invitrogen).

To generate stable cell lines, 68KD cells were seeded in 35 mm wells and co-transfected after 24 h with 0.3 μg of the appropriate pcDNA5/FRT-derived plasmid (empty/wild type CFIm68/mutant CFIm68) and 1.7 μg of pOG44 (Invitrogen) using JetPrime (Polyplus-transfection). Immediately before transfection, the medium was replaced with medium lacking zeocin and blasticidin. After 48 h, cells were passaged into medium supplemented with 15 μg/ml blasticidin (InvivoGen) and 150 μg/ml hygromycin B (InvivoGen) in a 10 cm dish to select for stable integrants. After appearance of colonies, cells were batch passaged and maintained in the blasticidin/hygromycin-supplemented medium.

### Mutagenesis

CFIm68 deletion and point mutants (other than RS1 S-to-A and ΔRS2, see below) were generated by PCR-based site-directed mutagenesis from the pcDNA5/FRT-CFIm68 backbone using Turbo Pfu polymerase (Stratagene) and complimentary oligonucleotides containing the desired mutation/deletion and flanking sequence.

To generate the RS1 S-to-A multi-site mutant, a series of oligonucleotides containing the desired mutations (CFIm68 RS1 S-to-A 1-5) were annealed and ligated together. The final construct with *Ava*I/*Pac*I sticky ends was ligated into the pcDNA5/FRT-CFIm68 plasmid following *Ava*I/*Pac*I restriction digestion.

To generate the ΔRS2 mutant, the ΔRS2 F and R oligonucleotides containing *Kpn*I and *Xho*I restriction sites were used to amplify the appropriate portion of the *CPSF6* cDNA lacking the C terminus from an existing plasmid, and the fragment was cloned into pcDNA5/FRT by restriction digestion (as for the wild type cDNA, see above).

### Western blotting, immunoprecipitation and phosphorylation analysis

To prepare lysates, cell pellets were resuspended in lysis buffer (50 mM Tris-HCl pH 7.8, 150 mM NaCl, 0.5% IgePal) supplemented with cOmplete EDTA-free protease inhibitors (Roche) and 10 mM benzamidine. For immunoprecipitation experiments, the buffer was further supplemented with PhosSTOP phosphatase inhibitor (Roche), 2.5 mM MgCl_2_ and 125 U/ml benzonase. Lysates were cleared by centrifugation at 16 100 x g for 18 min, and protein concentrations measured using Bradford reagent (BioRad).

For immunoprecipitation, 500-700 μg lysate was incubated with 4 μg antibody per mg lysate (CFIm68 IP) or 2.5 μg antibody per mg lysate (CFIm25 IP), followed by addition of protein G-Sepharose beads (Sigma) blocked in 3 mg/ml BSA. Following three washes with wash buffer (50mM Tris-HCl, 150 mM NaCl, 1 mM MgCl_2_, 0.05% IgePal + inhibitors), beads were boiled in 4 X LDS buffer (NuPage) before loading on the gel. For western blotting of whole cell lysates, 25-40 μg was mixed with 10 μl 4 X LDS buffer and boiled.

Samples (lysate/IP) were separated by SDS-PAGE (11% polyacrylamide) and transferred to a nitrocellulose membrane. For standard western blots, membranes were blocked in 5% milk in TBS-T before probing with indicated primary antibodies and IRDye 680-conjugated anti-rabbit IgG or IRDye 800CW-conjugated anti-mouse IgG secondary antibodies (LI-COR). For phosphorylation analysis, blocking buffers and detection reagents from the pIMAGO kit (Tymora Analytical), including the avidin-fluor 800 secondary, were used.

Blots were imaged and quantified using the Odyssey SA platform and software (LI-COR), using manual band drawing and the median background subtraction method.

### Primary antibodies

The following antibodies were used: CFIm25 western blotting - NUDT21 10322-1-AP (rabbit polyclonal, ProteinTech). CFIm25 immunoprecipitation – NUDT21 2203C3 (mouse monoclonal, sc-81109, SantaCruz). CFIm68 western blotting – CPSF68 A301-356A (rabbit polyclonal, Bethyl Laboratories). CFIm68 immunoprecipitation – CPSF68 A301-358A (rabbit polyclonal, Bethyl Laboratories). GFP immunoprecipitation – GFP B-2 (mouse monoclonal, sc-9996, SantaCruz). α-tubulin western blotting - Anti-α-tubulin, clone DM1A (mouse monoclonal, T9026, Sigma). GAPDH western blotting – Anti-GAPDH (rabbit polyclonal, G9545, Sigma). Normal rabbit IgG control immunoprecipitation – normal rabbit IgG (sc-2027, SantaCruz).

### Northern blotting

Total RNA was extracted by re-suspension of cell pellets in TRI reagent (Sigma) followed by bromochloropropane phase separation and isopropanol precipitation. 15-20 μg of RNA was denatured in loading buffer (50% formamide, 6% formaldehyde, 1x MOPS, 10% glycerol, 20 μg/ml ethidium bromide, 0.05% bromophenol blue) and separated on a 1.2% agarose-formaldehyde gel. RNA was transferred to a Hybond-N+ membrane (GE Healthcare) overnight by capillary action using 10x SSC, then fixed by UV crosslinking. RNA integrity was confirmed by ethidium bromide visualisation of 28S and 18S rRNA bands.

The membrane was pre-hybridised in hybridisation buffer (50% formamide, 10% dextran sulphate, 5x SSC, 1% SDS, 1x Denhardt’s solution, 100 μg/ml denatured salmon sperm DNA) before adding radiolabelled DNA probes for hybridisation at 42°C overnight.

Probes were generated by PCR from HEK 293 Flp-In T-Rex cDNA (with TIMP2/SDC2/ACTB oligonucleotides) and gel purified, then radiolabelled with [α-^32^P] dCTP using the DECAprime II DNA labelling kit (Ambion) with random priming.

After hybridisation, the membrane was washed in 1x SSC, 0.1% SDS for 15 min at 42°c followed by 15 min at 65°c, then twice in 0.1x SSC, 0.1% SDS at 65°c. Signals were visualised using a phosphor-imager, and bands were quantified by 2D densitometry using AIDA image analyser (Raytest).

### Statistical analysis

Statistical significance of differences between proximal/distal band ratios on northern blots was determined using a two-tailed, unpaired Student’s t-test. On graphs, significance level is represented by asterisks as follows: * p ≤0.05, ** p ≤0.01, *** p ≤0.001, **** p ≤0.0001. Error bars on graphs represent ± standard error of the mean (S.E.M.).

### GFP tagging and live cell confocal microscopy

Plasmids encoding N-terminal GFP fusions of CFIm68 were constructed by insertion of a GFP cDNA as a *Kpn*I fragment into pcDNA5/FRT-derived plasmids encoding wild type/ΔRS/ΔpNLS CFIm68.

Cells were imaged at RT on an Olympus FV1000 IX81 confocal microscope system using a 60x 1.35NA oil immersion lens and Olympus Fluoview software (Olympus, Southend-on-Sea, Essex, UK). Hoechst 33342 (1μg/ml) was added directly to the medium 15 minutes before imaging.

### Chromatin immunoprecipitation (ChIP) and deep sequencing analysis

ChIP was performed as previously described (52). Briefly, 293 Flp-In and 68KD cells were cross-linked at room temperature with 1% formaldehyde and quenched with 125 mM glycine for 5 minutes. Nuclear extracts were sonicated twice for 15 minutes at high amplitude, 30s ON/30s OFF using a Bioruptor (Diagenode). 80 μg of chromatin was incubated overnight at 4°C with 2 μg of an antibody against IgG (sc-2027, Santa Cruz) as an IP negative control or against pol II (sc-899X, Santa Cruz). After recovery of immune complexes with BSA-saturated protein G Dynabeads and extensive washes, crosslinks were reversed by incubation at 65°C for 5 hours. After ethanol precipitation and proteinase K treatment, DNA was purified using Qiagen PCR Purification Kit. ChIP samples were analysed by deep sequencing using Illumina HiSeq 4000 75 bp paired-end reads (Wellcome Trust Centre for Human Genetics, University of Oxford).

To analyse data, adapters were trimmed with Cutadapt v. 1.9.1 (53) with the following constant parameters: --minimum-length 10 −q 15, 10 –-max-n 1. Obtained sequences were mapped to the human hg19 reference sequence with Bowtie2 v. 2.2.5 (54). Unmapped reads were removed with SAMtools v. 1.3.1 (55). Mapped reads were then de-duplicated using Picard to remove PCR duplicates. Bam files were sorted and indexed with SAMtools. Data were normalized to Reads Per Genomic Content (RPGC) by employing deepTools2 v. 2.2.4 (56) bamCoverage tool with the following parameters: -bs 10 -normalizeTo1x 2451960000 -e –p max.

For retrieving non-overlapping transcription start sites (TSSs), the GENCODE V19 annotation was parsed using a custom Python script to keep only non-overlapping TSSs within a region of 2.5 kb upstream or downstream of the TSS. The metaprofile was created with deepTools2 v. 2.2.4 computeMatrix tool. ChIP-seq datasets have been deposited into the GEO genomics data repository (accession number: GSE99955).

### Oligonucleotides

Not1-CFIm68 F: GGGGAGCGGCCGCATGGCGGACGGCGTG

CFIm68-XhoI R: GGGGCTCGAGCTAACGATGACGATATTCGCGCTC

CFIm68 target site F: TGAGGGGGAAAATATCTTGCAGT

CFIm68 target site R: CCTCCTTCCAATGTAAACAATCATG

M13 R: CAGGAAACAGCTATGAC

CFIm68 Δ116-122 F: TGGAGATAAAATTTTTTTCAAAGGGGTTTGCCCTTG

CFIm68 Δ116-122 R: CAAGGGCAAACCCCTTTGAAAAAAATTTTATCTCCA

CFIm68 ΔRS1 F: GATTGCCTTCATGGAATTCGACGTCATAAATCCCGT

CFIm68 ΔRS1 R: ACGGGATTTATGACGTCGAATTCCATGAAGGCAATC

KpnI-CFIm68 ΔRS2 F: CGCGGTACCAGCGGCCGCATGGC

CFIm68 ΔRS2-XhoI R: CCGCCTCGAGCTATTTATGACGTCGACTCTTTTCTCG

CFIm68 ΔpNLS F: CCATAGTAGATCACGAGAAAAGAGTTCCCGTAGTAGAG

CFIm68 ΔpNLS R: CTCTACTACGGGAACTCTTTTCTCGTGATCTACTATGG

CFIm68 S511A F: TCGACGTCATAAAGCCCGTAGTAGAGACCGTC

CFIm68 S511A R: GACGGTCTCTACTACGGGCTTTATGACGTCGA

CFIm68 S513A F: CGTCATAAATCCCGTGCTAGAGACCGTCATGAC

CFIm68 S513A R: GTCATGACGGTCTCTAGCACGGGATTTATGACG

CFIm68 S511A S513A F: CGACGTCATAAAGCCCGTGCTAGAGACCGTCATGAC

CFIm68 S511A S513A R: GTCATGACGGTCTCTAGCACGGGCTTTATGACGTCG

CFIm68 S525A F: ATTACAGAGAGAGAGCCAGAGAACGAGAGAGG

CFIm68 S525A R: CCTCTCTCGTTCTCTGGCTCTCTCTCTGTAAT

CFIm68 S511D F: CGACGTCATAAAGACCGTAGTAGAGACCGTCATGA

CFIm68 S511D R: TCATGACGGTCTCTACTACGGTCTTTATGACGTCG

CFIm68 S513D (from WT) F: CGTCATAAATCCCGTGATAGAGACCGTCATGAC

CFIm68 S513D (from WT) R: GTCATGACGGTCTCTATCACGGGATTTATGACG

CFIm68 S513D (from S511D) F: CGTCATAAAGACCGTGATAGAGACCGTCATGAC

CFIm68 S513D (from S511D) R: GTCATGACGGTCTCTATCACGGTCTTTATGACG

CFIm68 S525D F: ATTACAGAGAGAGAGACAGAGAACGAGAGAGG

CFIm68 S525D R: CCTCTCTCGTTCTCTGTCTCTCTCTCTGTAAT

CFIm68 RS1 S-to-A 1 F:

TAAACAATCCAAAGTATCTGCTGATGATCGTTGCAAAGTTCTTATTA

GTTCTTT

CFIm68 RS1 S-to-A 1 R:

TGAAGGCAATCTTGCAAAGAACTAATAAGAACTTTGCAACGATCAT

CAGCAGATACTTTGGATTGTTTAAT

CFIm68 RS1 S-to-A 2 F:

GCAAGATTGCCTTCATGGAATTGAGGCCAAGGCTTATGGTGCTGGA

GCAAGACGTGAACGAG

CFIm68 RS1 S-to-A 2 R:

GGTCCCTCTCTCTTGCTCGTTCACGTCTTGCTCCAGCACCATAAGCC

TTGGCCTCAATTCCA

CFIm68 RS1 S-to-A 3 F:

CAAGAGAGAGGGACCATGCTAGAGCACGAGAAAAGGCTCGACGTC

ATAAATCCCGTAGTAGA

CFIm68 RS1 S-to-A 3 R:

ATCGTCATGACGGTCTCTACTACGGGATTTATGACGTCGAGCCTTTT CTCGTGCTCTAGCAT

CFIm68 RS1 S-to-A 4 F:

GACCGTCATGACGATTATTACAGAGAGAGAAGCAGAGAACGAGAG

AGGCACCGGGATCGTGA

CFIm68 RS1 S-to-A 4 R:

CGGTCACGGTCTCGGTCACGATCCCGGTGCCTCTCTCGTTCTCTGCT TCTCTCTCTGTAATA

CFIm68 RS1 S-to-A 5 F:

CCGAGACCGTGACCGAGAGCGTGACCGAGAGCGCGAATATCGTCAT

CGTTAGC

CFIm68 RS1 S-to-A 5 R: TCGAGCTAACGATGACGATATTCGCGCTCTCGGTCACGCTCT

TIMP2 F: CGCAACAGGCGTTTTGCAAT

TIMP2 R: TGGTGCCCGTTGATGTTCTT

SDC2 F: TGTACCTTGACAACAGCTCC

SDC2 R: GCCAATAACTCCACCAGCAA

ACTB F: GGATTCCTATGTGGGCGACG

ACTB R: GTAGTCAGTCAGGTCCCGGC

## Acknowledgments

We thank Andrew Bassett for the design, cloning and initial testing of the *CPSF6* sgRNA and Alan Wainman for valuable assistance with the GFP microscopy. This work was supported by Cancer Research UK (CR-UK) grant number C38302/A13012, through an Oxford Cancer Research Centre Prize DPhil Studentship.

## Author contributions

CJN conceived and designed the study. JGH designed and performed the experiments and analysed the data, except the RNA polymerase II ChIP-seq. SM conceived and MT performed the ChIP-seq experiments jointly with JGH and carried out bioinformatic analysis of the data. JGH and CJN jointly wrote the paper and all authors approved the final version of the manuscript.

